# Low humidity enhances Zika virus infection and dissemination in *Aedes aegypti* mosquitoes

**DOI:** 10.1101/2024.01.17.576075

**Authors:** Angel Elma I. Abu, Margaret Becker, Anastasia Accoti, Massamba Sylla, Laura B. Dickson

## Abstract

As climate change alters Earth’s biomes, it is expected the transmission dynamics of mosquito-borne viruses will change. While the effects of temperature changes on mosquito-virus interactions and spread of the pathogens have been elucidated over the last decade, the effects of relative humidity changes are still relatively unknown. To overcome this knowledge gap, we exposed *Ae. aegypti* females to various low humidity conditions and measured different components of vectorial capacity such as survival, blood-feeding rates, and changes in infection and dissemination of Zika virus. Survival decreased as the humidity level decreased, while infection rates increased as the humidity level decreased. Alternatively, blood feeding rates and dissemination rates peaked at the intermediate humidity level, but returned to the levels of the control at the lowest humidity treatment. These results provide empirical evidence that *Ae. aegypti* exposure to low humidity can enhance Zika virus infection in the mosquito, which has important implications in predicting how climate change will impact mosquito-borne viruses.

## Importance

Arthropod-borne viruses (arboviruses) such as Zika (ZIKV), dengue (DENV), and chikungunya (CHIKV) viruses transmitted by the mosquito, *Aedes aegypti*, pose a major threat to public health [1, 2]. Climate models predict that as global temperatures rise, the risk for arboviral disease will increase and that arboviruses will become a greater threat globally, and specifically in Africa [3, 4]. Global expansion of Zika [5, 6] and year-round transmission potential from *Ae. aegypti* is likely to expand particularly in South Asia and sub-Saharan Africa [3, 7]. While most predictions about the impact of climate change on vector-borne disease transmission are focused on temperature, little is known about how other climate variables will impact vector-borne disease dynamics. A climate variable tightly linked to temperature is humidity [8]. Given the importance of dehydration in mosquito physiology [9, 10], the variability in humidity across environments, and the predicted changes in humidity under climate change, it is imperative that we also study the impact that it has on mosquito infection and transmission of arboviruses.

## Observation

Water loss, or dehydration, has been linked to phenotypes impacting vectorial capacity. Water loss has been directly associated with changes in mosquito behavior, leading to an increase in blood feeding rate and potential increases in transmission of West Nile virus (WNV) [11-13]. Additionally, water loss has been connected to decreased survival and oviposition [14].

A single study has empirically measured the effect of low humidity on infection [15]. In this study, *Ae. aegypti* were exposed to either 35% RH, 75% RH, or 80% RH (control) for 18 hours prior to the infectious bloodmeal containing Mayaro virus (MAYV). Manzano-Alvarez et al. reported that there were no significant differences in infection, dissemination, or transmission rates (rate of MAYV infection in saliva) at either 7 dpi or 14 dpi between the treatments, and there was also no significant difference in survival. However, they did observe that the blood-feeding rate of the 75% group was significantly higher than the 35% group and control, which were similar to each other.

To confirm the impact of low humidity on various phenotypes related to vectorial capacity, *Ae. aegypti* collected from the city of Thíes in Senegal [16] were exposed to Zika virus under three relative humidity ranges, 20%, 50%, or 80% (control) for 24 hours prior to the infectious bloodmeal and for the duration of infection. We measured survival, blood-feeding, infection, and dissemination rates, as well as dissemination titers, to assess vector competence (Figure 1A). Detection of ZIKV RNA and infectious particles was done as previously described [17].

**Figure 1:**
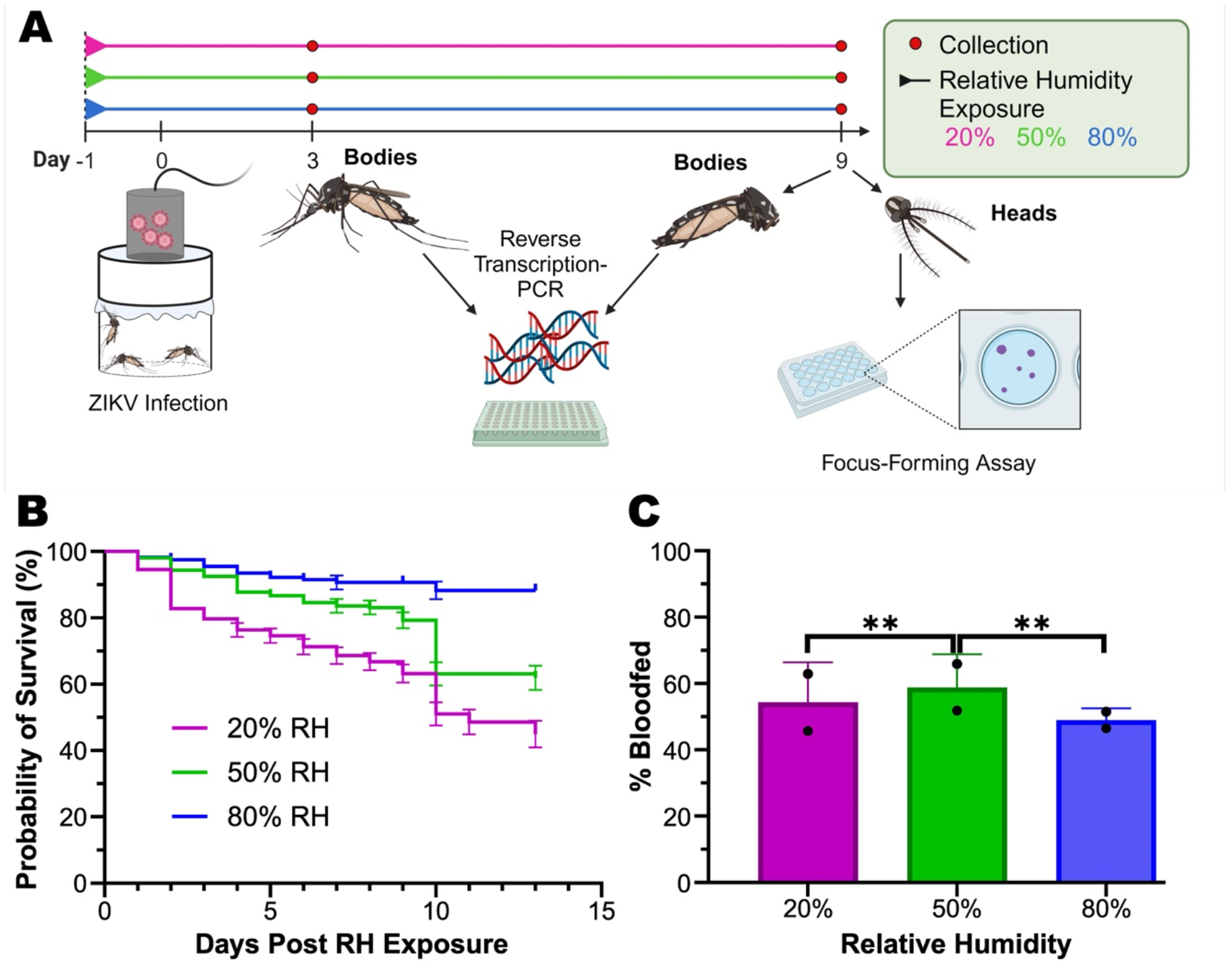
Low humidity impacts survival probability and blood-feeding rate. A) F12 *Aedes aegypti* collected in Thíes, Senegal [16] were exposed to 20% or 50% with a control of 80% relative humidity treatments 24 hours prior to the infectious blood feeding and maintained at those humidity levels for the duration of the experiment. Experiments were conducted in two biological replicates. On Day 0, mosquitoes were fed a bloodmeal infected with Zika virus (Cambodia FSS 13025 isolate) at a titer of 3.4 × 10^7^ FFU/mL. Collections were made at days three and nine to determine the infection and dissemination rates. Infection rates were determined in bodies by RT-PCR of the bodies three and nine days post infection (dpi), and dissemination rates and titers were determined in heads nine dpi through a focus forming assay as described previously [17]. B) Probability of survival over a span of 13 days post infectious bloodmeal for each treatment. A Log-Rank (Mantel-Cox) test was performed as a mean of two replicates (p-value < 0.0001) with sample sizes of 433 individuals from the 20% treatment, 427 from the 50% treatment, and 289 from the control. C) Blood-feeding rates were determined by the number of fully engorged mosquitoes divided by the total number that was offered the bloodmeal. The results are presented as a mean of two replicates from sample sizes of 869 (20%), 758 (50%), and 588 (80%). Overall and treatment vs. treatment significance was determined by a two-tailed Chi-squared analysis. P-values: Overall = 0.009752, 20% vs. 50% = 0.0071, 20% vs. 80% = 0.8495, 50% vs. 80% = 0.0088.

The relative humidity and temperature were monitored by digital hygrometers for the duration of the study. Following exposure to low humidity, mosquitoes exhibited lower survival probabilities as the humidity decreased (Figure 1B). Blood-feeding rate was also observed as a function of humidity. The 20% 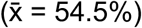and 80%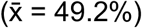humidity treatments did not have significantly different blood-feeding rates (p-value = 0.8495). However, they had lower mean feeding rates compared to the 50% humidity treatment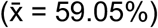,which was statistically significant when compared to the 20% (p-value = 0.0071) and 80% treatments (p-value = 0.0088) (Figure 1C). These results were similar to the previous study on MAYV in which they also observed an increase in blood-feeding rates in their intermediate humidity level [15].

To understand the impact that relative humidity has on ZIKV infection in *Ae. aegypti*, adult females were harvested at three- and nine-days post infection. While there were no significant differences in infection rates between three and nine days, mosquitoes maintained at 20% (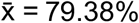, p-value = 0.0001) and 50% humidity (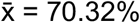, p-value = 0.0004) had significant increases in infection rates compared to the control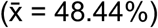. However, there was no significant difference between the 20% and 50% treatments (p-value = 0.1235) even though infection rates trended higher in the 20% treatment (Figure 2A).

**Figure 2:**
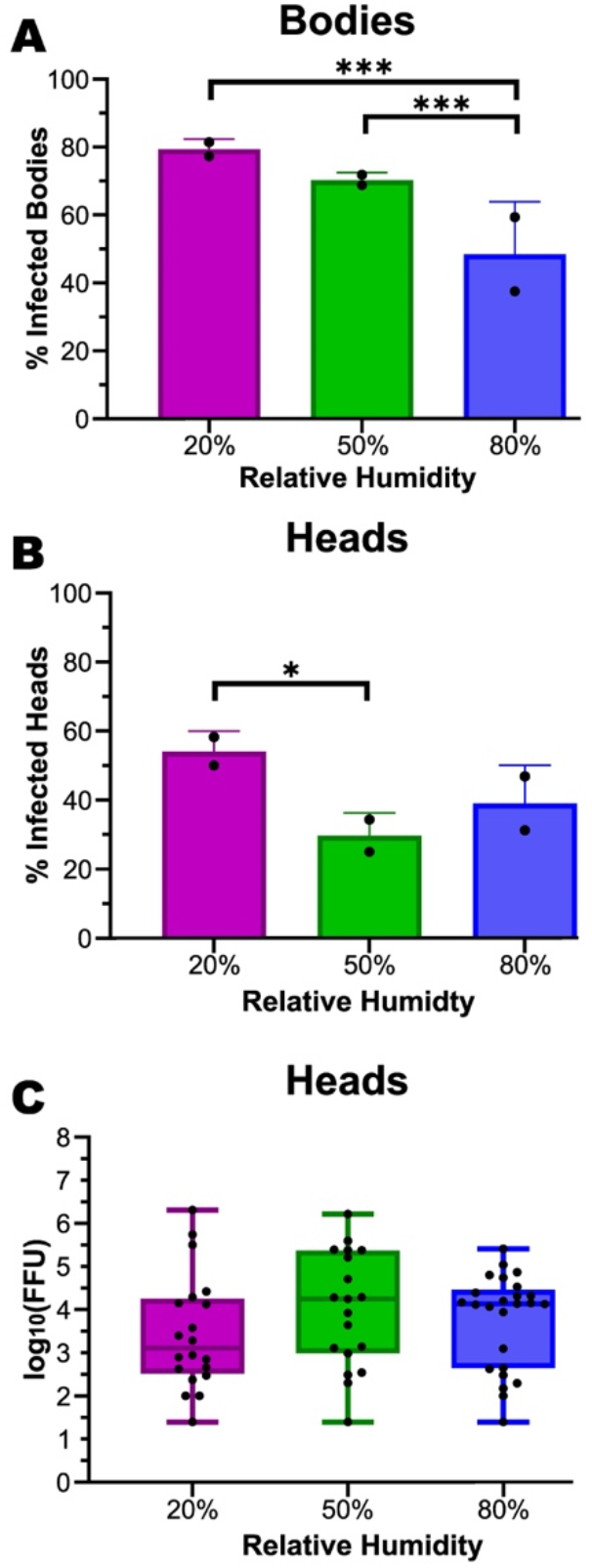
Low humidity increases infection and dissemination rates but has no impact on dissemination titers. A) The percentage of infected bodies is shown for each humidity treatment. Results from individuals collected three and nine dpi were combined due to there being no statistically significant difference between the two collection timepoints. Infection rates were determined by detecting ZIKV RNA by RT-PCR. A Log-Rank (Mantel-Cox) test was performed as a mean of two replicates (p-value = 1.93×10^6^) with sample sizes of 54 (20% RH), 64 (50% RH), and 64 (80% RH). Significance between treatments was assessed through a two-tailed Chi-square analysis with the following p-values: 20% vs. 50% = 0.1235, 20% vs. 80% = 0.0001, 50% vs. 80% = 0.0004. B) The percentage of infected heads is shown for each humidity treatment. Infection was determined in heads nine dpi through a focus forming assay. The percentage of infected heads was determined by the number of infected samples divided by the total number tested per treatment. Values were taken as the mean of two replicates. A Log-Rank (Mantel-Cox) test was performed (p-value = 0.03996) with sample sizes of 36 individuals from the 20% treatment, 64 from the 50% treatment, and 64 from the control. Significance between treatments was assessed through a two-tailed Chi-square analysis with the following p-values: 20% vs. 50% = 0.0109, 20% vs. 80%= 0.1115, 50% vs. 80% = 0.2642. C) Mean viral titer was calculated as the log10(FFU)/ml. All values were not significant. Significance was determined by ANOVA, p-value 0.4377, and two-tailed Chi-squared tests with the following p-values: 20% vs. 50% = 0.3878, 20% vs. 80% = 0.1777, 50% vs. 80% = 0.3843.

To test whether low humidity increases ZIKV dissemination, the proportion of mosquitoes that developed a disseminated infection were measured by detecting infectious virus particles in the head nine days post infection. Mosquitoes maintained at 20% (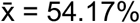, p-value = 0.1115) and 50% (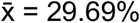, p-value = 0.2642) showed no significant difference in dissemination compared to the control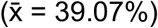. However, there were significant differences between the 20% and 50% treatments (p-value = 0.0071) (Figure 2B) with dissemination peaking at the 50% humidity level. Additionally, the quantity of infectious virus particles was measured in the heads to determine if low humidity exposure increased replication of the virus. The results showed a mean dissemination titer in log10(FFU) of 3.50 in the 20%, 4.01 in the 50%, and 3.76 in the 80%. Therefore, there were no significant differences in the amount of infectious virus particles in the heads between the treatments and control (p-value 0.4377) (Figure 2C).

In contrast to Manzano-Alvarez et al. [15] we observed that mosquito infection and dissemination rates increase in response to low humidity. Perhaps conflicting results between this study and Manzano-Alvarez et al. are a result of the different genetic backgrounds of *Ae. aegypti* used, different impacts of low humidity on alphaviruses versus flaviviruses, or differences between the timing and duration of low humidity exposure. The highly inbred Liverpool line of *Ae. aegypti* was used in the Manzano-Alvarez et al study, while a F12 line of *Ae. aegypti* collected in Thíes, Senegal [16]. Given the contribution of mosquito genetics to vector competence [18] and the involvement of a desiccation induced gene in midgut infection [19], it is possible that low humidity impacts differently across genetic backgrounds. Additionally, in Manzano-Alvarez et al, mosquitoes were exposed to their humidity treatments for only 24 hours post infectious bloodmeal, while in this study the humidity treatments were maintained for the duration of viral incubation.

Interestingly, an increase in dissemination rates was only observed at the intermediate humidity level, but not at the lowest humidity level. Additionally, blood-feeding rates returned to the same levels as the control in the lowest humidity treatment and only increased in the intermediate humidity treatment. This data suggests that the mechanisms underlying the influence of low humidity on virus dissemination and blood feeding rates are likely complicated and perhaps different mechanisms exist for different humidity ranges.

Overall, this study demonstrates that exposure to low humidity increases ZIKV infection and dissemination rates in *Ae. aegypti*, which has important implications for the effect of climate on the transmission of arboviruses. This experimental framework paves the way for future mechanistic studies on how variation in humidity influences ZIKV infection and replication in the mosquito.

## Acknowledgements

We would like to thank Jiehua Zhou and Ruimei Yun of the UTMB Insectary Core. We would like to thank Dr. Noah Rose and Dr. Caroline McBride for sharing the mosquito colony with us. LBD was supported by UTMB start-up funds and NIH grant U01AI151801, the West African Center for Emerging Infectious Diseases. MB was a recipient of the T32 predoctoral fellowship. NIAID T32 Emerging and Tropical Infectious Diseases Training Program (AI007526 to Lynn Soong). This research was also supported by NIH grant R24 AI120942.

AEIA and LBD conceived the study. AEIA, MB, AA, and LBD performed the work. AEIA and LBD analyzed the data. MS provided the mosquito line. AEIA and LBD wrote the paper.

